# A Gallium-68-Labeled Peptide Radiotracer For CD38-Targeted Imaging In Multiple Myeloma With PET

**DOI:** 10.1101/2023.05.09.540036

**Authors:** Ajay Kumar Sharma, Kuldeep Gupta, Akhilesh Mishra, Gabriela Lofland, Ian Marsh, Dhiraj Kumar, Gabriel Ghiaur, Philip Imus, Steven P. Rowe, Robert F. Hobbs, Christian B. Gocke, Sridhar Nimmagadda

**Affiliations:** The Russell H. Morgan Department of Radiology and Radiological Science, Johns Hopkins University School of Medicine, Baltimore, MD, 21287, USA; Chemical & Biomolecular Engineering, Whiting School of Engineering, Johns Hopkins University School of Medicine, Baltimore, MD, 21287, USA; The Sidney Kimmel Comprehensive Cancer Center and the Bloomberg–Kimmel Institute for Cancer Immunotherapy, Johns Hopkins University School of Medicine, Baltimore, MD, 21287, USA; Department of Pharmacology and Molecular Sciences, Johns Hopkins University School of Medicine, Baltimore, MD, 21287, USA; Division of Clinical Pharmacology, Department of Medicine, Johns Hopkins University School of Medicine, Baltimore, MD, 21287, USA

**Keywords:** CD38, PET, Daratumumab, Isatuximab, ATRA, Minimal residual disease

## Abstract

**Purpose:** The limited availability of molecularly targeted low-molecular-weight imaging agents for monitoring multiple myeloma (MM)-targeted therapies has been a significant challenge in the field. In response, we developed [^68^Ga]Ga-AJ206, a peptide-based radiotracer that can be seamlessly integrated into the standard clinical workflow and is specifically designed to non-invasively quantify CD38 levels and pharmacodynamics by positron emission tomography (PET).

**Experimental design:** We synthesized a high-affinity binder for quantification of CD38 levels. Affinity was tested using surface plasmon resonance, and *In vitro* specificity was evaluated using a gallium-68-labeled analog. Distribution, pharmacokinetics, and CD38 specificity of the radiotracer were assessed in MM cell lines and in primary patient-derived myeloma cells and xenografts (PDX) with cross-validation by flow cytometry and immunohistochemistry. Furthermore, we investigated the radiotracer’s potential to quantify CD38 pharmacodynamics induced by all-trans retinoic acid therapy (ATRA).

**Results:** [^68^Ga]Ga-AJ206 exhibited high CD38 binding specificity (K_D_: 19.1±0.99 nM) and CD38-dependent *In vitro* binding. [^68^Ga]Ga-AJ206-PET showed high contrast within 60 minutes and suitable absorbed dose estimates for clinical use. Additionally, [^68^Ga]Ga-AJ206 detected CD38 expression in xenografts, PDXs and disseminated disease models in a manner consistent with flow cytometry and immunohistochemistry findings. Moreover, [^68^Ga]Ga-AJ206-PET successfully quantified CD38 pharmacodynamics in PDXs, revealing increased CD38 expression in the tumor following ATRA therapy.

**Conclusions:** [^68^Ga]Ga-AJ206 exhibited the salient features required for clinical translation, providing CD38-specific high contrast images in multiple models of MM. [^68^Ga]Ga-AJ206-PET could be useful for quantifying total CD38 levels and pharmacodynamics during therapy to evaluate approved and new therapies in MM and other diseases with CD38 involvement.

**STATEMENT OF TRANSLATIONAL RELEVANCE:** There is an unmet need for functional imaging agents to monitor the pharmacodynamic effects of new therapeutics targeting multiple myeloma (MM). MM is a challenging bone marrow plasma cell cancer that is associated with heterogenous responses and universal recurrence. While minimal residual disease monitoring by blood and invasive bone marrow samples have improved prognostication of disease recurrence, molecularly targeted, non-invasive imaging options that can assess therapy response early remain limited. To address this gap, we report the development of a high affinity, first-in-class gallium-68 labeled peptide radiotracer, [^68^Ga]Ga-AJ206, for CD38 protein, which is highly expressed on MM cells. [^68^Ga]Ga-AJ206 provides high-contrast CD38-specific images by PET within the standard clinical workflow of 60 minutes. Furthermore, the potential of [^68^Ga]Ga-AJ206 PET to measure pharmacodynamics of CD38 was demonstrated. Further development of this new radiotracer may complement existing technologies and improve prognostication and monitoring of therapy response in patients with MM.

## INTRODUCTION

Multiple myeloma (MM) is a plasma cells neoplasm that primarily arises in the bone marrow (1). Clinically, MM is characterized by wide variability in response to therapy and survival thought to be due to biologic heterogeneity in both cell-intrinsic and -extrinsic factors (1). Therapeutic options for MM include combinations of drugs with different mechanisms of action, such as proteasome inhibitors (PIs), immunomodulatory drugs (IMIDs), B cells maturation antigen (BCMA) targeting agents, monoclonal antibodies targeting MM antigens (mAbs, including bispecific monoclonal antibodies (2), chimeric antigen receptor T cells (CAR-T), and high-dose chemotherapy rescued by autologous stem cell transplantation (ASCT). Current methods to quantify the burden of MM in patients includes bone marrow biopsies, serum testing of monoclonal paraprotein, and cross-sectional imaging (3,4). Among the latter, positron emission tomography combined with computed tomography (PET-CT) with the metabolic radiotracer 2-deoxy-2-[^18^F]fluoro-D-glucose ([^18^F]FDG) is the best equipped to determine response to treatment and provide prognostic information (5). The significant limitations of [^18^F]FDG include lack of specificity due to inflammatory processes such as healing fractures. However, some cases of myeloma are not FDG-avid(6). The development of molecularly targeted PET imaging agents that enhance MM detection specificity, sensitivity, and early therapeutic response assessment remains an unmet need.

Cluster of differentiation protein 38 (CD38) is expressed uniformly and with high density in MM. Taking advantage of high CD38 expression, several therapies including antibodies, antibody drug conjugates and chimeric antigen receptor-T (CAR-T) cells have been developed or are in different phases of clinical development (7). Continuous therapy with a combination of the aforementioned agents has been associated with the best outcomes in MM (8). However, there are efforts underway to reduce toxicity by tailoring or tapering treatment in patients who have little or no detectable minimal residual disease (MRD) (9). Also, efforts are underway to increase CD38 expression on MM cells to take advantage of the excellent therapeutic options available (10,11). As a result, accurate detection of small amounts of disease will become increasingly important for this growing patient population. Realizing the need to improve specificity and sensitivity to detect MM, several groups have leveraged the high expression of CD38 on MM cells (12-21) to develop molecularly targeted imaging agents. Radiolabeled conjugates of anti-CD38 antibodies have been shown to detect MM in preclinical models and patients, with superior specificity and sensitivity compared to the routinely used [^18^F]FDG-PET (12,18). However, the long half-life of those agents limits their applicability to patient who have not received anti-CD38 therapy. Additionally, although the diagnostic and therapeutic implications are less-well understood, CD38 is expressed in other cancers, including prostate cancer, and molecularly targeted agents may have a more pan-cancer role in the long-term.

Peptides are gaining traction as effective tools for targeting cancer due to their high specificity, rapid clearance, and easy synthesis (22). In particular, peptide-radiotracers are utilized to improve contrast and enhance the detection of cancerous lesions (23). Compared to antibodies, peptides have shorter clearance times (hours instead of days), making them a favorable option for repeated imaging for assessing response to therapies. The tractable pharmacokinetic properties of peptide-based imaging agents also enable the quantification of target pharmacodynamics (24,25). Notably, the application of low molecular weight agents for targeting CD38 in multiple myeloma (MM) has not been previously reported. In this study, we capitalized on these characteristics to develop a first-in-class peptide-based low molecular weight agent that allows for the quantification of CD38 levels and pharmacodynamics throughout the body within a few hours.

Here, we report the development and evaluation of a new gallium-68 labeled peptide-based radiotracer, [^68^Ga]Ga-AJ206, for CD38 targeting in multiple myeloma (MM). We assessed the pharmacokinetics, biodistribution, and CD38 specificity of [^68^Ga]Ga-AJ206 *In vitro* and *In vivo* using multiple MM cell lines with variable CD38 levels, primary MM cells and their xenografts, and in disseminated tumor models. Additionally, we investigated the potential of [^68^Ga]Ga-AJ206-PET to detect the pharmacodynamics of CD38 during all-trans retinoic acid (ATRA) therapy.

## RESULTS

### Synthesis and characterization of AJ206

CD38-binding low-molecular-weight imaging agents have not been reported and high-affinity peptide therapeutics (IC_50_<100 nM) were reported only recently, which form the basis for our imaging agents **(Figure 1**)(26). To test the applicability of those peptides for *In vivo* imaging, we selected a peptide with high hydrophilicity and a free amine for further functionalization. We synthesized a *2,2*′,*2”-(1,4,7-triazacyclononane-1,4,7-triyl)triacetic acid* (NOTA)-conjugated peptide, AJ206, with a poly ethylene glycol (PEG) linker to improve solubility (**Figure 1A, and supplementary schemes S1 & S2**). The agent was characterized by mass spectrometry (**Supplementary figure 1)**. We then tested its potential to bind human (hCD38) and mouse CD38 (mCD38) recombinant proteins and MM cells with variable CD38 expression. AJ206 exhibited high affinity for hCD38 with a dissociation constant (K_D_) of 19.1 nM by Surface Plasmon Resonance (SPR) but did not bind mCD38 (**Figure 1B and supplementary figure S2**).

**Figure 1.**
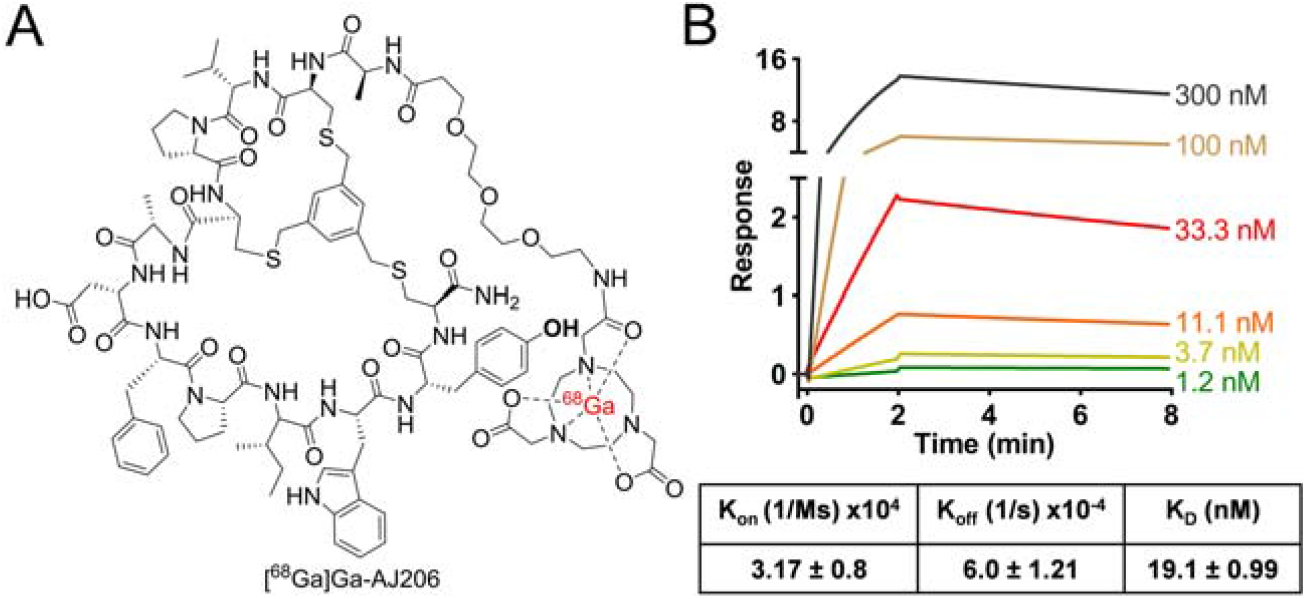
Structure and *In vitro* characterization of AJ206. **A)** Structure of bicyclic peptide AJ206 having NOTA as bifunctional chelator for ^68^Ga-labeling **B)** Surface plasmon resonance (SPR) analysis showing affinity of AJ206 for CD38 using recombinant human CD38 protein; data is represented as mean±SEM (n=2).

### Pharmacokinetics of [^68^Ga]Ga-AJ206

We synthesized the radioactive analog of AJ206, [^68^Ga]Ga-AJ206, by standard ^68^Ga-radiolabeling chemistry (**Supplementary scheme 3)** to evaluate the pharmacokinetics, biodistribution, and *In vivo* specificity (27). [^68^Ga]Ga-AJ206 was obtained with decay corrected radiochemical yields of 92±10.5% (n=35) with >95% radiochemical purity and a specific activity (molar activity) of 10-15 GBq/μmol (270-400 mCi/μmol) (**Supplementary figure 3)**. The formulated [^68^Ga]Ga-AJ206 dose remained stable for 2 h (**Supplementary figure 4**). Using immunodeficient NSG mice with MOLP8 human MM xenografts that are known to express CD38 (28), we evaluated [^68^Ga]Ga-AJ206 *In vivo*. Whole body dynamic PET/MR images of those mice acquired over 90 min showed clear and high radiotracer accumulation in tumors as early as few minutes after injection, with a higher tumor contrast observed at 60 min due to radiotracer clearance (**Figure 2A**). Kidney and bladder tissues had the highest radiotracer accumulation among normal tissues, consistent with the renal clearance mechanism for low-molecular-weight peptides, while radioactivity from other tissues, including liver, declined over 60 min (**Figure 2B**). These imaging data showed that [^68^Ga]Ga-AJ206 exhibits *In vivo* kinetics and contrast desirable for an imaging agent. To verify the PET results, we conducted *ex vivo* biodistribution studies (**Table 1 and Supplementary table 1**). At 60 min, [^68^Ga]Ga-AJ206 uptake in MOLP8 tumors peaked at 3.76±0.25% %ID/g (**Figure 2C)**. Blood pool activity exhibited consistent washout, with almost 95% of activity cleared by 60 min, and comparable radioactivity washouts were observed in all other non-specific tissues. As a result, the tumor-to-blood and tumor-to-muscle ratios were highest between 60 and 120 min (**Figure 2D, Table 1**). Thus, we chose the 60-minute time point for all further experiments because it provided good contrast and aligned with the standard PET imaging time frame.

**Table 1.** Kinetics and distribution of [^68^Ga]Ga-AJ206 in mice with MOLP8 tumor xenografts; data is presented as mean±SEM (n=4 or 5) of %ID/g.

| Tissues | 5 min | 30 min | 60 min | 120 min |
| --- | --- | --- | --- | --- |
| Blood | 8.25±0.50 | 2.07±0.30 | 0.56±0.09 | 0.05±0.01 |
| Muscle | 0.92±0.11 | 0.37±0.05 | 0.14±0.02 | 0.07±0.04 |
| Tumor | 2.58±0.18 | 2.91±0.84 | 3.76±0.25 | 1.46±0.02 |
| Heart | 3.13±0.27 | 0.83±0.18 | 0.29±0.03 | 0.05±0.01 |
| Lungs | 6.16±0.16 | 1.86±0.29 | 0.71±0.10 | 0.16±0.03 |
| Liver | 21.24±1.31 | 6.39±3.71 | 1.22±0.09 | 0.42±0.01 |
| Spleen | 2.28±0.50 | 0.75±0.14 | 0.39±0.03 | 0.17±0.02 |
| Kidney | 11.17±0.65 | 17.09±1.77 | 16.65±2.52 | 13.75±0.23 |
| Stomach | 1.08±0.07 | 0.51±0.06 | 0.25±0.06 | 0.03±0.01 |
| Small intestine | 1.95±0.08 | 0.59±0.14 | 0.65±0.09 | 0.18±0.04 |
| Femur | 1.42±0.09 | 0.65±0.08 | 0.28±0.07 | 0.07±0.01 |
| Tumor/Muscle | 2.94±0.47 | 7.28±1.89 | 27.12±2.73 | 27.37±18.55 |
| Tumor/Blood | 0.32±0.04 | 1.43±0.46 | 7.20±0.89 | 19.72±6.57 |

**Figure 2.**
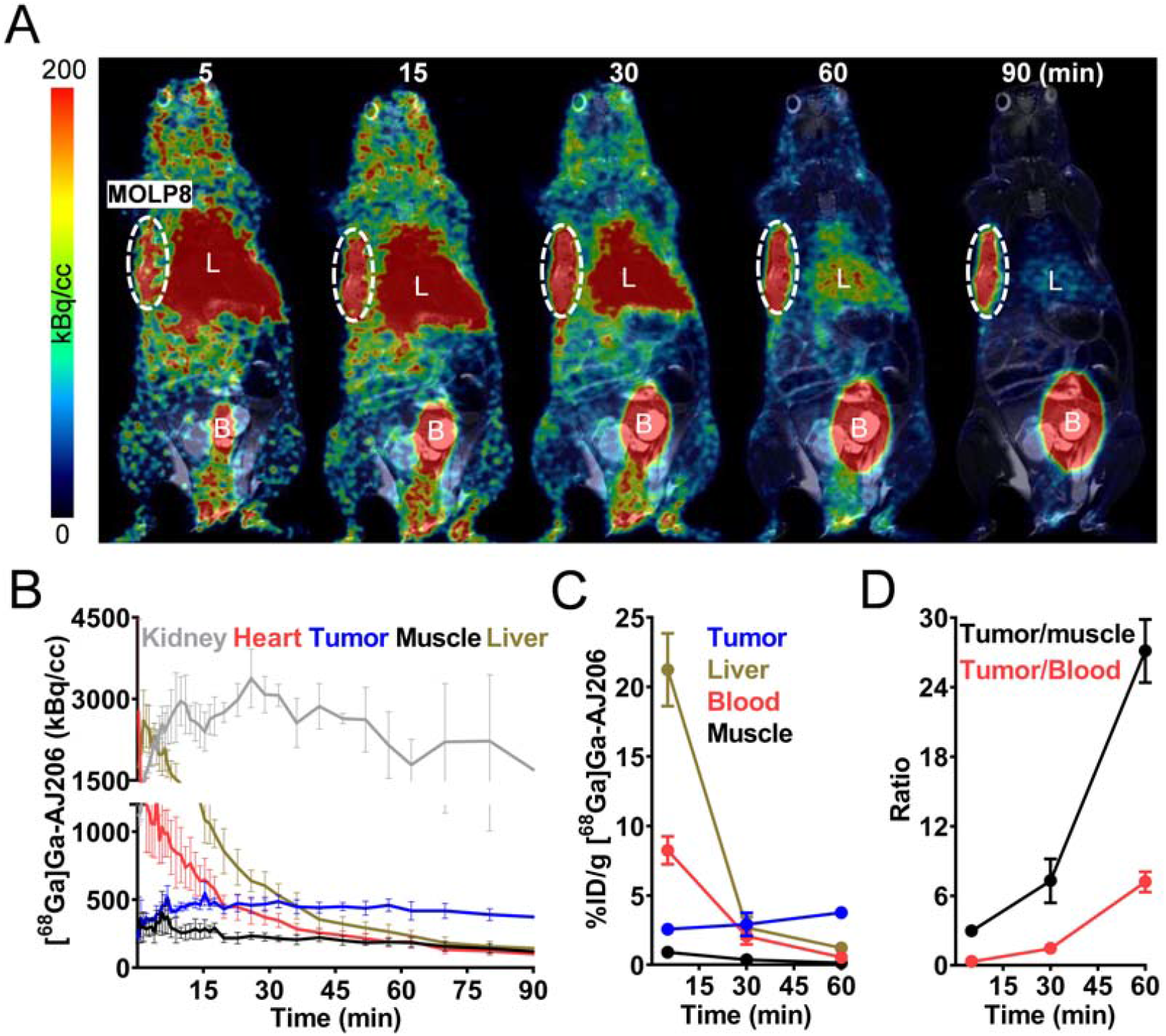
Pharmacokinetics of [^68^Ga]Ga-AJ206 in NSG mice bearing MOLP8 tumor xenografts. **A)** Coronal sections of the fused dynamic PET/MR images showing [^68^Ga]Ga-AJ206 distribution. Primary tumor is indicated by the dashed white circle. Mice were intravenously injected with ∼ 9.25 MBq (∼250 μCi) [^68^Ga]Ga-AJ206; L, Liver; B, Bladder. **B)** Time-activity curves of [^68^Ga]Ga-AJ206 in the kidney, heart, liver, tumor and muscle derived from images A. **C)** Uptake of [^68^Ga]Ga-AJ206 in tumor, blood, liver and muscle derived from *ex vivo* biodistribution study. **D)** Tumor-to-muscle and tumor-to-blood ratios derived from biodistribution data. Data in panels **C** and **D** are from mice intravenously injected with ∼ 2.96 MBq (∼80 μCi) [^68^Ga]Ga-AJ206 and sacrificed at different time-points after injection, represented as mean±SEM (n=4 or 5).

**Figure 3.**
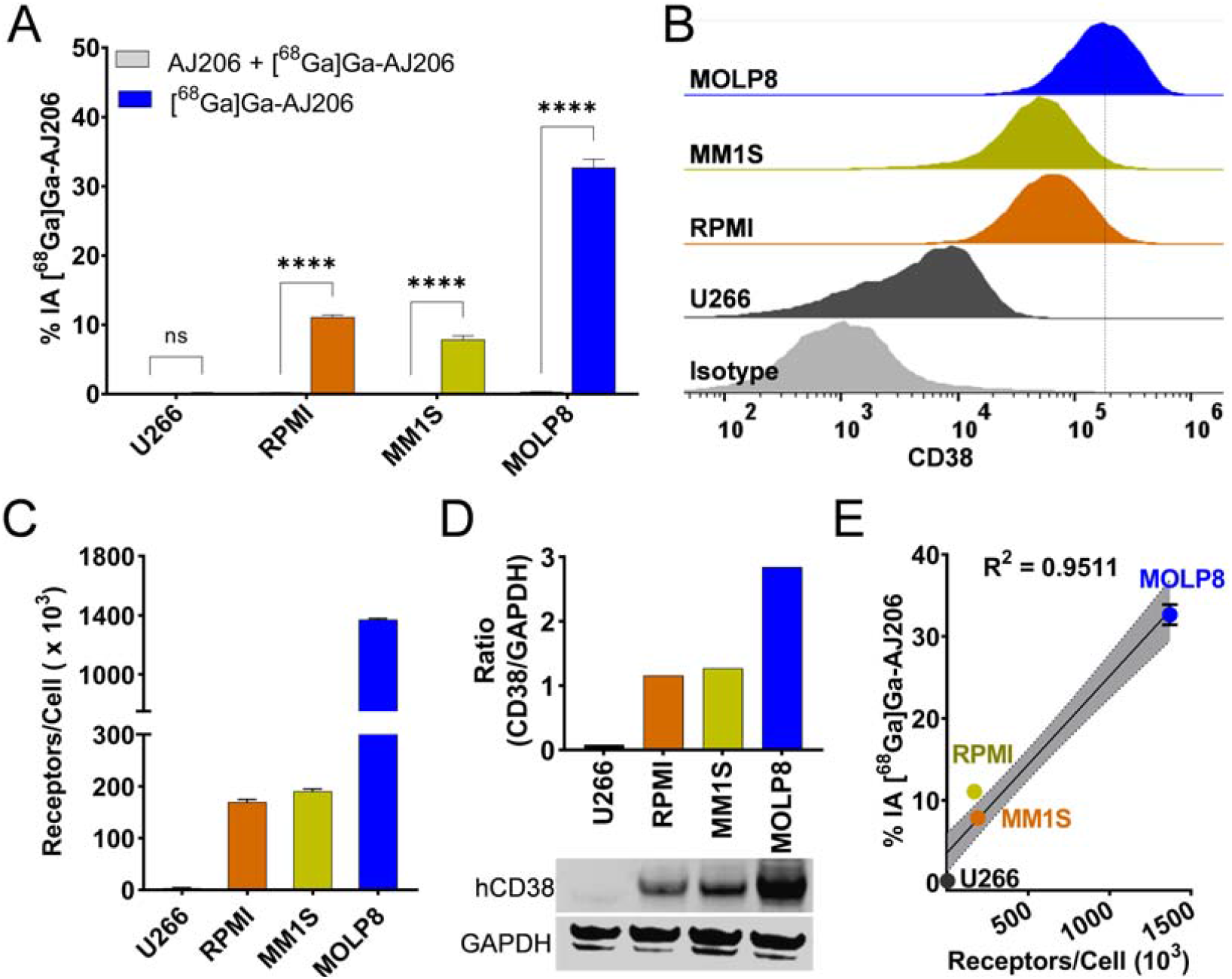
*In vitro* specificity of [^68^Ga]Ga-AJ206 for CD38. **A)** [^68^Ga]Ga-AJ206 binding (percent incubated activity, %IA) to different MM cells. Cells were incubated with 1 μCi [^68^Ga]Ga-AJ206 at 4 °C for 1 hour. [^68^Ga]Ga-AJ206 uptake is CD38 expression dependent, and co-incubation with 2 μM of non-radioactive AJ206 (blocking dose) significantly reduced radiotracer uptake confirming CD38 specificity. **B)** Flow cytometry analysis of CD38 surface expression in MM cells. **C)** CD38 receptor density in MM cells measured by quantibrite assay. **D)** Representative western blot of total CD38 protein expression (bottom panel). Densiometric analysis of western blot preformed using ImageJ software and band intensities represented as a ratio of CD38 protein to GAPDH control (top panel). **E)** Correlation of [^68^Ga]Ga-AJ206 uptake with surface CD38 receptor density; data in panels **A, C** and **E** are represented as mean±SD (n=3-4). ns, *P* ≥ 0.05; ****, *P* ≤ 0.0001 is by unpaired Student’s *t test*. Simple linear regression and Pearson coefficient were used in **E**.

**Figure 4.**
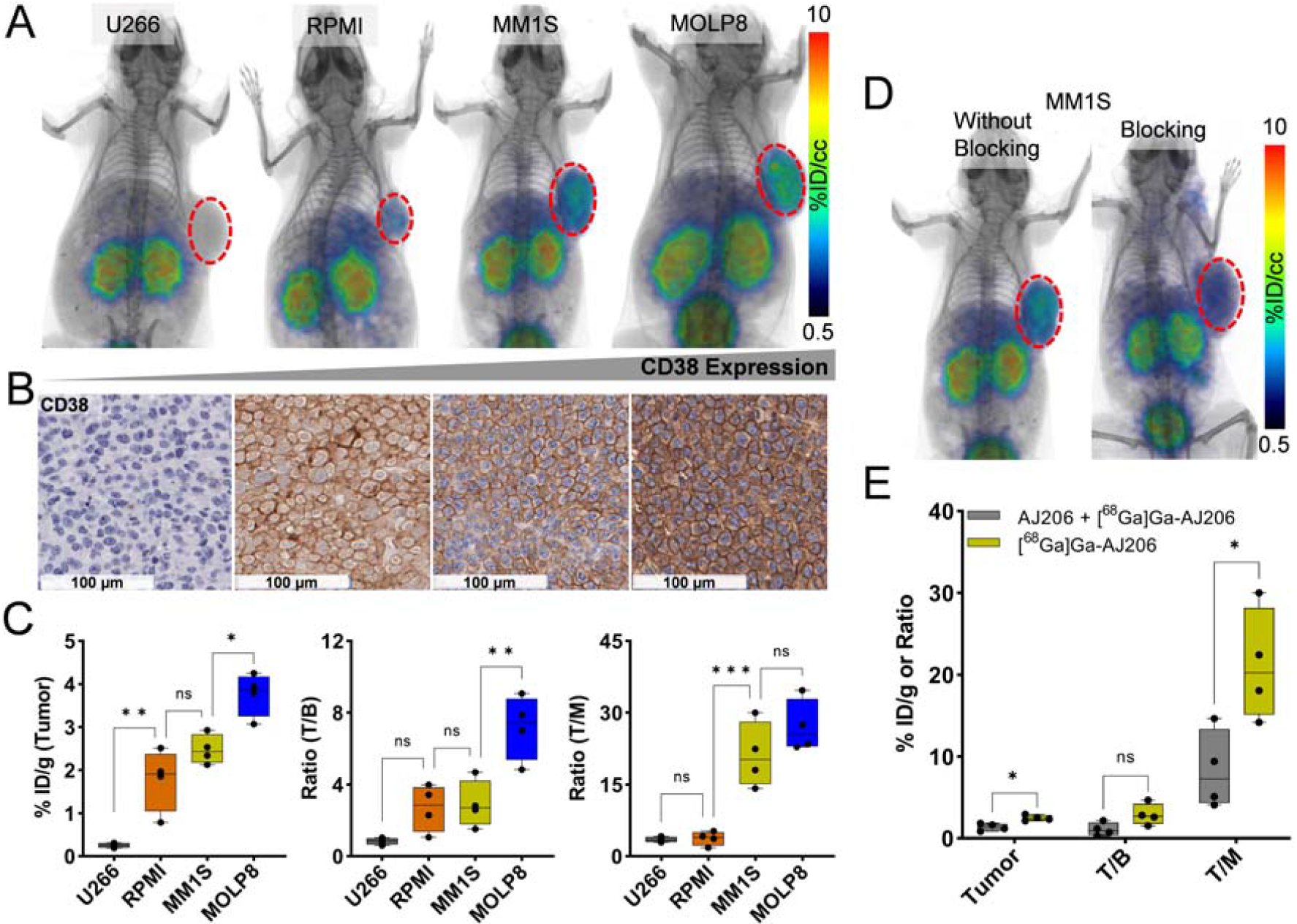
In vivo specificity of [^68^Ga]Ga-AJ206 for CD38 in NSG mice with MM tumor xenografts. **A)** Whole-body PET/CT images of different human MM xenografts at 60 min after the injection of radiotracer. Mice were injected with ∼ 7.4 MBq (∼200 μCi) [^68^Ga]Ga-AJ206. **B)** IHC staining for CD38 expression in MM xenografts. **C)** [^68^Ga]Ga-AJ206 uptake quantification (%ID/g) in different MM tumors by ex vivo biodistribution at 60 min after injection. **D)** PET/CT images of MM1S tumor xenograft-bearing mice with [^68^Ga]Ga-AJ206, with and without pre-administration of a blocking dose (50 μg of AJ206) (tumor denoted with dashed red line). **E)** [^68^Ga]Ga-AJ206 quantification in tumors by ex vivo biodistribution in mice treated with and without a blocking dose; data in figure **C**, and **E** are shown as box and whisker plots showing all data points (n=4-5). Ordinary one-way ANOVA using multiple comparison test in **C** and multiple unpaired *t* test in **E**. ns, P ≥ 0.05; *, P ≤ 0.05; ** P ≤ 0.01; ***, P ≤ 0.001.

### Human Radiation Dosimetry Estimates

Pharmacokinetic data obtained from biodistribution studies of MM1S tumor-bearing mice was used to predict the time-integrated activity coefficients (TIACs, previously known as residence times) of [^68^Ga]Ga-AJ206 in humans (**Supplementary table 1 and 2**). TIAC estimates were then employed as input to MIRDCalc to calculate the absorbed dose coefficients to organs from ^68^Ga using the ICRP adult female reference phantom (**Supplementary table 2**). The kidneys received the highest absorbed dose (0.13 rem/mCi), followed by the liver (0.06 rem/mCi), heart wall (0.05 rem/mCi), and lung (0.05 rem/mCi). Based on these results, a 30 mCi dosage can be safely administered, with an estimated effective dose equivalent of less than 5 rem, to obtain PET images.

### Confirmation of [^68^Ga]Ga-AJ206 specificity in MM cell lines and xenografts with varying CD38 levels

Next, we conducted an evaluation of the specificity of [^68^Ga]Ga-AJ206 to accurately detect varying levels of CD38 expression both *In vitro* and *In vivo*. Four MM cell lines (U266, RPMI, MM1S, and MOLP8) were incubated with [^68^Ga]Ga-AJ206, and the amount of cell-bound radioactivity was measured. We observed that MOLP8 and U266 cells showed the highest and the least amount of radioactivity binding, respectively, while RPMI and MM1S cells demonstrated intermediate uptake (**Figure 3A**). We confirmed the specificity of [^68^Ga]Ga-AJ206 binding by observing a significant reduction in radiotracer uptake (*P* < 0.0001) when cells were incubated with a saturating concentration (2 μM) of non-radioactive AJ206. To validate the *In vitro* uptake, we performed flow cytometry analysis to measure cell surface expression and density, and western blot analysis to assess the total CD38 protein expression. Our results showed that the highest cell surface receptor expression and density were observed in MOLP8 cells followed by RPMI and MM1S cells, with the lowest expression in U266 cells (**Figure 3B and 3C**). Similarly, MOLP8 cells exhibited the highest total CD38 expression, followed by RPMI and MM1S cells, while U266 cells had the lowest (**Figure 3D**). We also performed a correlation analysis to investigate the relationship between [^68^Ga]Ga-AJ206 uptake and CD38 receptor density, and found a strong correlation (*R*^*2*^ = 0.9511), indicating that [^68^Ga]Ga-AJ206 uptake in cells is CD38 specific (**Figure 3E**).

To validate the *In vitro* results, we performed PET imaging studies of MM xenografts derived from the cell lines mentioned above, using [^68^Ga]Ga-AJ206. PET images were acquired at 60 min and showed highest radioactivity accumulation in MOLP8 tumors, followed by MM1S and RPMI tumors, and least accumulation in U266 tumors (**Figure 4A**). To corroborate the PET data, we performed IHC staining of the same xenografts for CD38 and found intense staining in MOLP8 xenografts and least in U266 tumors (**Figure 4B**). Additionally, we conducted *ex vivo* biodistribution studies to further validate the imaging findings. We found that the tumor uptake was consistent with *In vitro* and PET results, with the highest uptake observed in MOLP8 tumors (3.76±0.25 %ID/g), followed by MM1S (2.48±0.17 %ID/g) and RPMI (1.7±0.36). On the other hand, U266 tumors showed 0.25±0.03 %ID/g of [^68^Ga]Ga-AJ206, which is in line with their low CD38 levels. The tumor-to-muscle and tumor-to-blood ratios showed similar trends with respect to CD38 status (**Figure 4C**), and non-specific tissues showed no significant differences between tumor models (**Supplementary table 3**). Furthermore, we demonstrated the specificity of [^68^Ga]Ga-AJ206 for CD38 by co-administering 2 mg/kg non-radioactive AJ206, which significantly reduced the radioactivity uptake in MM1S xenografts (*P* < 0.05) (**Figure 4D and 4E**). These *In vitro* and *In vivo* data together demonstrate the potential of [^68^Ga]Ga-AJ206 for non-invasive detection of varying levels of CD38 expression.

**Figure 4. *In vivo* specificity of [**^**68**^**Ga]Ga-AJ206 for CD38 in NSG mice with MM tumor xenografts. A)** Whole-body PET/CT images of different human MM xenografts at 60 min after the injection of radiotracer. Mice were injected with ∼ 7.4 MBq (∼200 μCi) [^68^Ga]Ga-AJ206. **B)** IHC staining for CD38 expression in MM xenografts. **C)** [^68^Ga]Ga-AJ206 uptake quantification (%ID/g) in different MM tumors by *ex vivo* biodistribution at 60 min after injection. **D)** PET/CT images of MM1S tumor xenograft-bearing mice with [^68^Ga]Ga-AJ206, with and without pre-administration of a blocking dose (50 μg of AJ206) (tumor denoted with dashed red line). **E)** [^68^Ga]Ga-AJ206 quantification in tumors by *ex vivo* biodistribution in mice treated with and without a blocking dose; data in figure **C**, and **E** are shown as box and whisker plots showing all data points (n=4-5). Ordinary one-way ANOVA using multiple comparison test in **C** and multiple unpaired *t* test in **E**. ns, *P* ≥ 0.05; *, *P* ≤ 0.05; ** *P* ≤ 0.01; ***, *P* ≤ 0.001.

### Validation of [^68^Ga]Ga-AJ206 specificity for CD38 in disseminated MM disease models and primary plasma cell leukemia xenografts

We next conducted a study to evaluate the potential of [^68^Ga]Ga-AJ206 to visualize disease in bone marrow and soft tissues for non-invasive monitoring of high-risk MM phenotypes. Extramedullary manifestation (EMD) and plasma cell leukemia (PCL) are two aggressive clinical presentations of MM that exhibit high relapse rates. EMD presents with infiltration in other organs, such as lymph nodes, liver, lungs, and central nervous system, while PCL is characterized by free circulation of MM cells in the blood. To achieve this, we generated a disseminated disease model by intravenously injecting luciferase expressing MM1S to track cell engraftment (**Figure 5A**) or MOLP8 cells into NSG mice. [^68^Ga]Ga-AJ206 PET showed specific accumulation of radioactivity in bone lesions and other organs in both MM1S (**Figure 5B**) and MOLP8 tumor models (**Figures 5D**). This finding was further validated by bioluminescence imaging in the MM1S model (Figure 5A) and *ex vivo* PET imaging in the MOLP8 model (Figure 5D). IHC analysis confirmed high expression of CD38 in mice injected with MM1S-Luc (**Figure 5C)** and MOLP8 cells **(Figure 5E)**. Moreover, analysis of bone lesions from MM1S-Luc and MOLP8 models showed differential [^68^Ga]Ga-AJ206 uptake, indicating that different CD38 levels can be differentiated by [^68^Ga]Ga-AJ206 PET (**Figures 5F and 5G**). We also extracted cells from the bone marrow of those mice, which showed high CD38 expression, thus confirming that the observed uptake is indeed CD38 specific (**Figure 5H**).

**Figure 5.**
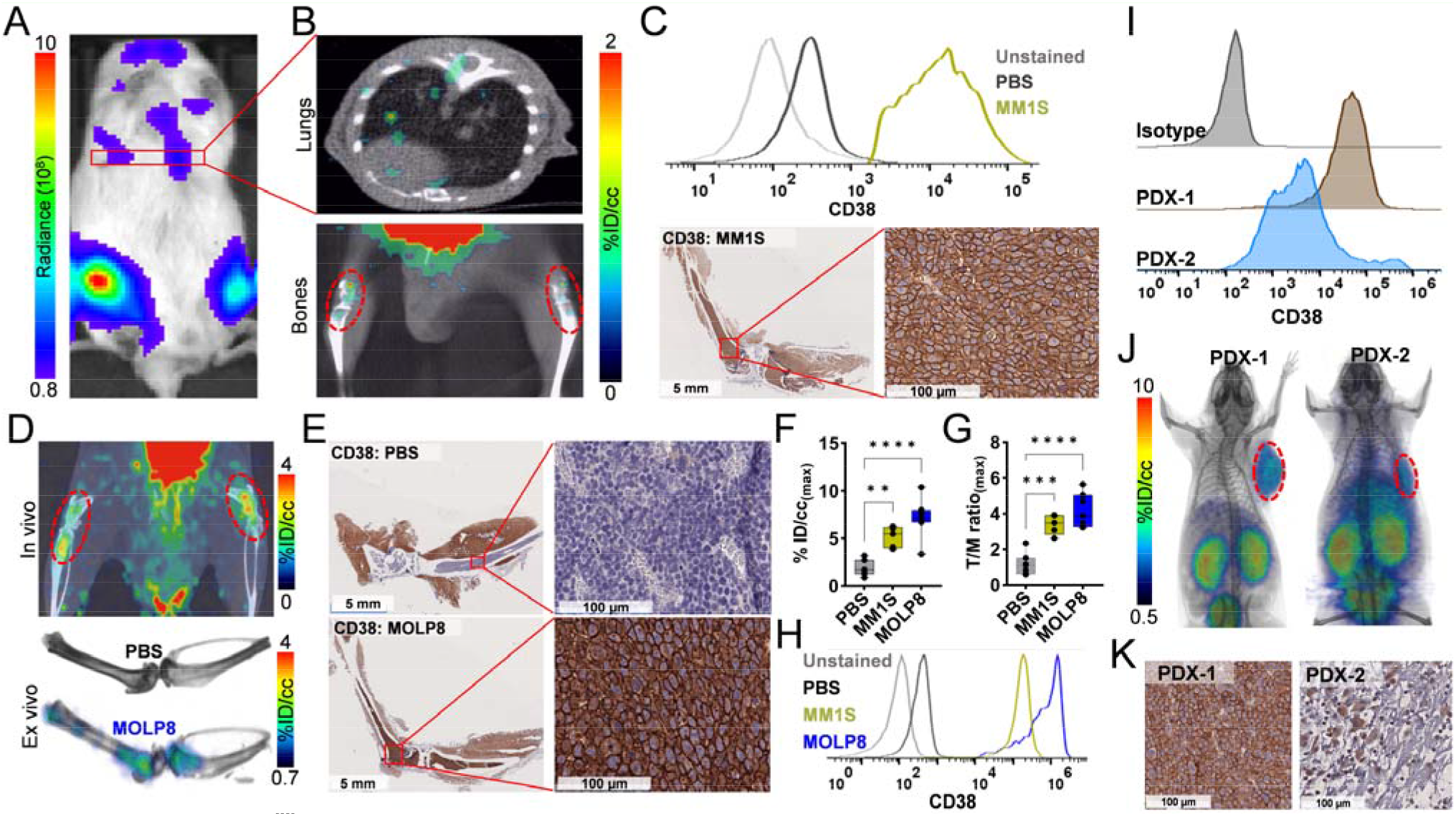
Validation of [^68^Ga]Ga-AJ206 specificity for CD38 in disseminated MM disease models and primary plasma cell leukemia xenografts. **A)** IVIS-bioluminescence image of luciferase expressing MM1S disseminated tumor model. **B)** *In vivo* uptake of [^68^Ga]Ga-AJ206 in lungs and bones in MM1S-Luc bearing mice. **C)** Flow cytometry analysis of CD38 expression in lungs harvested from MM1s-Luc cells injected mice. Lungs from PBS treated mice were used as controls (top panel). IHC staining of bones shows CD38 expression (bottom panel). **D)** *In vivo* and *ex vivo* [^68^Ga]Ga-AJ206-PET images of lower limbs of MOLP8 injected animals. **E)** IHC images of MOLP8 bone marrow tumor and PBS-treated animals confirmed CD38 expression. **F)** Quantification of PET signal (%ID/cc) in the bone marrow of PBS, MM1S-Luc and MOLP8 injected animals. **G)** Tumor/muscle (T/M) ratios of PET measures in the bone marrow of PBS, MM1S and MOLP8 injected animals. **H)** Flow cytometry analysis of CD38 expression in cells extracted from the bone marrow of PBS, MM1S-Luc and MOLP8 cell injected mice. **I)** Flow cytometry histograms of CD38 expression in primary cells. PDX-1 is from the peripheral blood of a relapsed/refractory MM patient with secondary plasma cell leukemia and PDX-2 is from a newly diagnosed MM patient bone marrow. Neither had exposure to anti-CD38 therapies. **J)** Static whole-body PET/CT images of PDX bearing mice at 60 min post injection of [^68^Ga]Ga-AJ206. (tumor in dashed red lines) **K)** IHC analysis of CD38 expression in PDXs. Data in figure F and G are shown as box and whisker plots showing all data points (n=7 in PBS, n=5 in MM1S and n=8 in MOLP8). Ordinary one-way ANOVA using multiple comparison test. **, *P* ≤ 0.01; *** *P* ≤ 0.001; ****, *P* ≤ 0.0001.

Extending our studies of [^68^Ga]Ga-AJ206 to PDXs, we first assessed the CD38 expression on MM cells from two anonymized patient samples and found expression within the range of previous reported expression levels (**Figure 5I)**. We then established primary patient-derived xenografts (PDXs) and carried out imaging studies. [^68^Ga]Ga-AJ206 PET showed specific accumulation of radioactivity in both the PDXs that reflected the CD38 expression detected by IHC **(Figures 5J and 5K)**. Collectively, these results indicate that [^68^Ga]Ga-AJ206 could aid in detection and monitoring of MM in both soft tissue and bone marrow.

### Pharmacodynamic Evaluation of CD38 with [^68^Ga]Ga-AJ206 in PDX Models

We next evaluated the potential of [^68^Ga]Ga-AJ206 to visualize therapy induced changes in CD38 levels. MM is often treated with the anti-CD38 antibody daratumumab, but its response is heterogeneous and linked to CD38 receptor density. Daratumumab treatment can also cause transient downregulation in CD38 expression and heterogenous responses(29). As a result, there are efforts underway to improve daratumumab efficacy by increasing CD38 expression using treatments such as ATRA (10,11). To investigate the potential of [^68^Ga]Ga-AJ206 to non-invasively evaluate therapy induced changes in CD38 expression, we first treated four MM cell lines with ATRA and measured changes in CD38 expression. Our results showed that ATRA treatment indeed increased CD38 expression (**Figure 6A**). Furthermore, when we incubated ATRA treated cells with [^68^Ga]Ga-AJ206, we observed a significant increase in radioactivity uptake compared to control treatment (**Figure 6B**). To validate the *In vitro* results, we performed PET imaging studies in PDX models. A significant increase in radioactivity uptake in tumors in mice were observed following treatment with ATRA compared to pre-treatment (**Figure 6C, D and supplementary figure 8A**). Also, [^68^Ga]Ga-AJ206 uptake in the PDXs was heterogeneous (**Figure 6D-F**), which correspond to variable CD38 expression detected in the flow cytometry and IHC analysis of those tumors (**Figure 6G and supplementary figure 8B, C**). In conclusion, this study provides evidence that [^68^Ga]Ga-AJ206 has the potential to detect therapy induced changes in CD38 expression *In vitro* and *In vivo*, which could be useful in efforts to improve the efficacy of anti-CD38 therapies.

**Figure 6.**
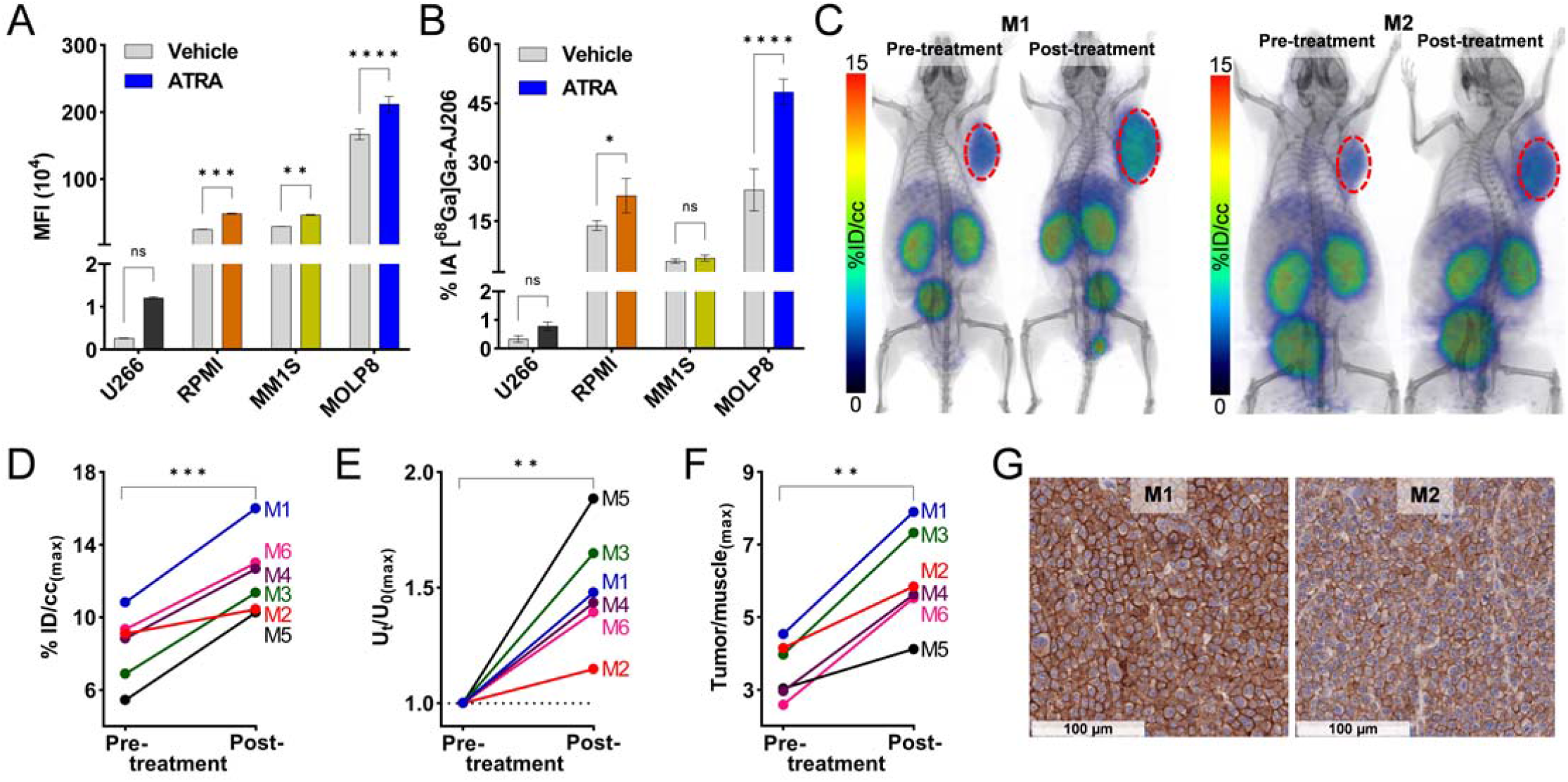
*In vitro* and *In vivo* detection of ATRA treatment induced changes in CD38 expression by [^68^Ga]Ga-AJ206 PET in MM cells and PDXs. **A)** Flow cytometry analysis of ATRA induced changes in surface expression of CD38 in MM cells. **B)** *In vitro* uptake of [^68^Ga]Ga-AJ206 (%IA) in MM cells treated with ATRA or vehicle control. Cells were incubated with [^68^Ga]Ga-AJ206 at 4 °C for 1 hour. **C)** Static whole-body PET/CT images of PDX bearing NSG mice before and after treatment with ATRA. Red circles indicate tumor. **D)** Quantification of PET signal in tumors pre- and post-treatment with ATRA. **E)** Ratio of PET signal in tumors before (U_0_) and after ATRA (U_t_) treatment. **F)** Tumor/Muscle ratio of PET measures before and after ATRA treatment **G)** CD38 IHC of PDXs from the same mice post-ATRA treatment; data in figures **A** and **B** are represented as mean±SD (n=3 or 4) and significance was calculated by ordinary two-way ANOVA using multiple comparison test; data in figure **D, E** and **F** are shown for individual mice and significance was calculated using paired *t test*. ns, *P* ≥ 0.05; *, *P* ≤ 0.05; **, *P* ≤ 0.01; ***, *P* ≤ 0.001; ****, *P* ≤ 0.0001.

## DISCUSSION

In this study, we demonstrated that a new ^68^Ga-labeled radiotracer, [^68^Ga]Ga-AJ206, exhibits high affinity for CD38 and optimal pharmacokinetics with excellent image contrast within 60 minutes after administration, aligning with standard clinical workflow and potentially allowing for facile translation into human imaging. Through tumor models, we confirmed that [^68^Ga]Ga-AJ206 detects varying levels of CD38 expression and differentiates CD38 levels in bone lesions in disseminated disease models. Our pre-clinical findings indicate that [^68^Ga]Ga-AJ206 has the potential to be a useful radiotracer in clinical settings for monitoring therapy response and disease progression in MM. Beyond MM, the expression of CD38 in other types of cancer, which is generally more heterogeneous than in MM, suggests that [^68^Ga]Ga-AJ206 may have a role in selecting those patients that might benefit from CD38 directed therapies. Across all types of CD38 expressing cancers, [^68^Ga]Ga-AJ206 provides an opportunity to identify new imaging biomarkers for disease prognosis.

Taking advantage of high CD38 expression in MM, several CD38-targeting therapies have been developed or are in different phases of clinical development (7). These include monoclonal antibodies such as daratumumab and isatuximab, which are approved for treating MM, as well as other forms of therapies such as ADCs, T-cell engagers, and CAR-T cells (7,30). Responses to anti-CD38 mAbs has been linked to CD38 receptor density and MM cells with higher levels of CD38 expression are more susceptible to anti-CD38-mediated therapeutic effects compared to low CD38 expression levels (7,10,29,31). Also, there is substantial heterogeneity in response among patients treated with these therapies (10,32-34). For example, daratumumab has been shown to induce durable responses in heavily pretreated patients, but the majority of responding patients eventually develop progressive disease during daratumumab monotherapy. Primary resistance to daratumumab has also been linked to CD38 receptor density (35). There is a current focus on enhancing CD38 expression on multiple myeloma (MM) cells in order to capitalize on the wide range of effective treatment options and improve overall efficacy (10,11). Consequently, the precise detection of minimal residual disease will gain greater significance for the expanding patient population, enabling them to benefit from emerging therapeutic alternatives. These clinical observations suggest that real-time, whole-body measurements of CD38 expression using molecularly targeted PET radiotracers could provide more accurate information for guiding anti-CD38 therapies in MM than current single-time point and single-lesion flow cytometry measurements. Moreover, non-invasive CD38 monitoring is expected to have a broader application across different disease states of MM, and may be applicable to a range of malignancies with known CD38 involvement, such as NK/T cell lymphoma, T-cell acute lymphoblastic leukemia, and primary effusion lymphoma.

Considering the importance of CD38 expression in MM, radiolabeled and fluorescence conjugates of daratumumab have been tested in preclinical models and patients. Several clinical trials are ongoing to capture the heterogeneity in CD38 levels in the whole body using radiolabeled analogs of the FDA approved anti-CD38 antibodies (mAbs). Radiolabeled anti-CD38 mAb (^89^Zr-daratumumab) localization to osseous MM sites was visualized by PET with appreciable contrast by day 7 (18). In another clinical trial using ^64^Cu-daratumumab, lesions that were ^18^F-FDG negative but positive in CD38-PET were found to be positive for MM involvement on biopsy (36), indicating the potential for CD38-PET for improved disease burden assessment. Our peptide tracers are small (∼2.1 kDa), synthetic, exhibit tractable pharmacokinetics, and take advantage of the inherent sensitivity of PET and the higher contrast resolution characteristics of ^68^Ga. As such, we observed the potential of [^68^Ga]Ga-AJ206 to detect varying CD38 levels in a variety of tumor models that is corroborated by CD38 IHC. Furthermore, our results suggest that CD38 specific low-molecular-weight diagnostic agents may have a number of applications in MM, including assessment of total CD38 levels, predicting response to therapy, and determining CD38 occupancy by mAbs, similar to what has been shown by us for PD-L1 therapeutics (24,25,37). Also, clinical trial evaluation of ATRA-induced changes in CD38 expression was evaluated in peripheral blood samples but not in bone marrow samples due to the invasive nature of the procedure (38). The potential of [^68^Ga]Ga-AJ206 in quantifying the pharmacodynamics of CD38 during therapy strongly suggests that non-invasive monitoring of CD38 levels may find use in how we monitor and assess the effectiveness of various therapies.

## MATERIALS AND METHODS

### Chemicals

NOTA-NHS ester was purchased from CheMatech, and all Fmoc protected amino acids, TIPS, HOBT, and HBTU were obtained from Chem-impex. Fmoc-PEG_3_-COOH and ^*t*^Bu_3_-DOTA-COOH were purchased from Ambeed, while DIPEA, TFA, DODT, TBMB, and DMF were obtained from Sigma-Aldrich.

### Synthesis of AJ205

AJ205 was chemically synthesized using Liberty Blue CEM automatic peptide synthesizer employing Fmoc based solid-phase peptide synthesis on Rink Amide resin in a 0.1 mmol scale (**Supplementary scheme 1**). Coupling reaction was carried out using Oxyma (0.5 mmol), DIC (1 mmol) and Fmoc-AA-OH (0.5 mmol) in DMF with microwave assisted reaction for 2 min. Fmoc group was deprotected using 20% piperidine in DMF (3 mL) for 1 min with microwave assistance. Next, PEG linker was incorporated by using HBTU (0.5 mmol), HOBT (0.5 mmol), DIPEA (0.5 mmol) and Fmoc-NH-PEG_3_-COOH (0.5 mmol) in DMF at room temperature for 1.5 h and then Fmoc group was deprotected using 20% piperidine (4 mL) in DMF for 40 min at room temperature. Once the sequence was completed on the resin, the peptidyl-resin was treated with 4 mL of cleavage cocktail (TFA:TIPS:DODT:H_2_O; 92.5:2.5:2.5:2.5) for 4 h at room temperature. The cleaved reaction mixture was precipitated with diethyl ether to obtain the linear peptide as a white solid.

For the cyclization, the linear peptide (98 mg, 0.06 mmol) was dissolved in 100 mL of water:acetonitrile (1:1) and treated with an aqueous solution of tris(2-carboxyethyl)phosphine (TCEP) (115 mg, 0.4 mmol) followed by Et_3_N (400 μL, 2.9 mmol). 1,3,5-Tris(bromomethyl)benzene (TBMB) (34 mg, 0.1 mmol) was dissolved in 2 mL of acetonitrile and added slowly to the reaction mixture over an hour. The reaction mixture was stirred at room temperature for 24 h and quenched with TFA. The volatiles were removed, and the crude mixture was purified on a reversed-phase high-performance liquid chromatography (RP-HPLC) system using a preparative C-18 Phenomenex column (5 mm, 21.5 x 250 mm Phenomenex, Torrance, CA). The HPLC condition was gradient elution starting with 10% acetonitrile: water (0.1% TFA) and reaching 60% acetonitrile: water (0.1% TFA) in 40 min at a flow rate of 8 mL/min. The product AJ205 was collected at retention time (RT) ∼28.2 min. Acetonitrile was evaporated under reduced pressure and lyophilized to form an off-white powder with a 37% yield (40 mg). The sequence of the AJ205 peptide is NH_2_-PEG_3_-ACVPCADFPIWYC-NH_2_ with TBMB-based cyclization. This peptide was characterized by MALDI-TOF-MS. The theoretical chemical formula is C_87_H_118_N_16_O_20_S_3_ with an exact mass of 1802.79 and a molecular weight of 1804.17. The theoretical MALDI-TOF-MS mass [M + H]^+^ was 1803.8; the observed ESI-MS mass [M + H]^+^ was 1804.0.

### Synthesis of AJ206

To a stirred solution of AJ205 (2.1 mg, 1.2 μmoles) in 200 μL of DMF in a reaction vial, NOTA-NHS ester (2.0 mg, 2.8 μmoles) and DIPEA (5 μL, 25 μmoles) were added and stirred at room temperature for 3 h (**Supplementary scheme 2**). DMF was evaporated using a rotary evaporator under reduced pressure and the residual product was purified on a RP-HPLC system using a semi-preparative C-18 Luna column (5 mm, 10 x 250 mm Phenomenex, Torrance, CA). The HPLC condition was gradient elution started with 5% acetonitrile: water (0.1% TFA) and reached at 95% acetonitrile: water (0.1% TFA) in 20 min at a flow rate of 5 mL/min. The product AJ206 was collected at RT ∼11.8 min. Acetonitrile was evaporated under reduced pressure and lyophilized to form an off-white powder with a 52% yield, which was characterized by MALDI-TOF-MS. The theoretical chemical formula of AJ206 is C_99_H_137_N_19_O_25_S_3_ with an exact mass of 2087.92 and molecular weight of 2089.47. The theoretical MALDI-TOF-MS mass [M + H]^+^ is 2088.9, and the observed ESI-MS mass [M + H]^+^ is 2089.1.

### Affinity measurements by surface plasmon resonance (SPR)

The affinity of AJ206 for hCD38 and mCD38 recombinant proteins were evaluated by SPR. The experiments were conducted using a Biacore T200 instrument with a CM5 chip at 25 °C. The ligands used were His-Tagged human CD38 (R&D systems, catalog # 2404-AC, 43 kDa, 0.5 mg/ml stock concentration) and mouse CD38 proteins (R&D systems, catalog # 4947-AC, 40 kDa, 1.81 mg/ml stock concentration), which were immobilized onto the CM5 chip. AJ206 (2089.2 Da, 10 mM stock concentration) was used as the analyte, which flowed over the ligand immobilized surface. FC2 was used as the experimental flow cell, while FC1 served as the reference. Anti-His antibody (1 mg/ml stock concentration) was immobilized on both FC1 and FC2 using standard amine coupling chemistry. The immobilization running buffer used was PBS-P (20 mM phosphate buffer pH 7.4, 137 mM NaCl, 2.7 mM KCl, 0.05% v/v surfactant P20). Human CD38 was captured onto FC2 at a level of ∼860 RU, with a 1:20 dilution and 25 μg/ml diluted concentration in PBS-P. Mouse CD38 was captured onto FC4 at a level of 1360 RU, with a 1:50 dilution and 36.2 μg/ml diluted concentration in PBS-P. The theoretical R_max_ values were calculated based on the captured response values and are presented in table 1, assuming a 1:1 interaction mechanism. Overnight kinetics were performed for all analytes in the presence of PBS-P+1% DMSO. The flow rate of all analyte solutions was maintained at 50 μL/min. The contact and dissociation times used were 120s and 360s, respectively. Surface regeneration was achieved by injecting glycine pH 1.5 for 20 seconds, which takes away all captured ligands onto FC2. Fresh ligands were captured at the beginning of each injection cycle. The analyte concentrations injected ranged from 300 nM down to 1.2 nM with three-fold serial dilutions, and all analytes were injected in triplicate.

| Ligand | FC | Analyte | Ligand Binding (RU) | MWL (Da) | MWA(Da) | Stoichiometric Ratio | R<sub>max</sub> |
| --- | --- | --- | --- | --- | --- | --- | --- |
| Human CD38 | 2 | AJ206 | 860 | 40000 | 2089.6 | 1:1 | 39.2 |
| Mouse CD38 | 4 | AJ206 | 1360 | 40000 | 2089.6 | 1:1 | 71.0 |

### Synthesis of [^nat^Ga]Ga-AJ206

The synthesis of [^nat^Ga]Ga-AJ206 was carried out by adding 10 μL of aqueous 0.1M [^nat^Ga]GaCl3 solution and 0.6 mL of 0.1 M HCl to a stirred solution of AJ206 (0.1 mg, 0.05 μmoles) in 200 μL of 1M NaOAc buffer (pH 5.0) in a reaction vial. The reaction mixture was incubated at 65 °C for 30 min and then purified on a RP-HPLC system using a semi-preparative C-18 Luna column (5 mm, 10 x 250 mm Phenomenex, Torrance, CA). The HPLC gradient elution condition started with 5% acetonitrile: water (0.1% TFA) and reached at 95% acetonitrile: water (0.1% TFA) in 20 min at a flow rate of 5 mL/min. The product [^nat^Ga]Ga-AJ206 was collected at RT ∼11.8 min. The acetonitrile was evaporated under reduced pressure and lyophilized to form an off-white powder, which was characterized by MALDI-TOF-MS. The theoretical chemical formula is C_99_H_134_GaN_19_O_25_S_3_ with an exact mass of 2153.82 and a molecular weight of 2156.17. The theoretical MALDI-TOF-MS mass [M + H]^+^ was 2154.8, and the observed ESI-MS mass [M + H]^+^ was 2154.1.

### Synthesis of [^68^Ga]Ga-AJ206

The ^68^Ge/^68^Ga generator was manually eluted using 6 mL of 0.1M HCl (Ultrapure trace-metal-free) in four different fractions (2.4 mL, 1 mL, 1 mL, and 1.4 mL). To a microcentrifuge vial (1.5 mL) containing 200 μL of 1 M NaOAc buffer (pH = 5) and 20 μg of AJ206 (10 nmoles), 3-4 mCi of [^68^Ga]GaCl_3_ in 0.6 mL from the second fraction was added (**Supplementary scheme 3**). The reaction mixture was incubated for 12 min at 65 °C in a temperature-controlled heating block and purified on a RP-HPLC system using a semi-preparative C-18 Luna column (5 mm, 10 x 250 mm Phenomenex, Torrance, CA). The HPLC condition was gradient elution starting with 5% acetonitrile: water (0.1% TFA) and reaching 95% acetonitrile: water (0.1% TFA) in 20 min at a flow rate of 5 mL/min. The radiolabeled product [^68^Ga]Ga-AJ206 was collected at RT ∼11.8 min, with decay-corrected radiochemical yield of 92±10.5 % (n=35). The desired radiolabeled fraction was concentrated under a stream of N_2_ at 60 °C, formulated in 10% EtOH in saline, and used for *In vitro* and *In vivo* studies. The whole radiolabeling process was completed in approximately 35 min. Quality control, stability studies, and chemical identity were also performed on the same HPLC system using the same HPLC gradient as described above.

### Cell Culture

MOLP8 cells were purchased from Deutsche Sammlung von Mikroorganismen und Zellkulturen (DSMZ), Germany, under the material transfer agreement guidelines with Johns Hopkins University. Other cell lines (U266, RPMI8226, MM1S and MM1S-Luc) were gifted by a collaborator. Prior to the start of the experiments, all cell lines were authenticated by STR profiling at the Johns Hopkins Genetic Resources Facility. Routine Mycoplasma testing was conducted, and the cell lines were passaged for no more than three months. New cultures were initiated from vials of frozen cells. All cell lines (U266, RMPI8226, MM1S, MM1S-Luc, and MOLP8) were cultured in the recommended media and maintained in an incubator at 37 °C in an atmosphere containing 5% CO2. All cells were supplemented with fetal bovine serum, 1% P/S antibiotics, and 1mM L-Glutamine.

### *In vitro* cellular binding assays

*In vitro* binding assays were conducted to determine the binding of [^68^Ga]Ga-AJ206 to U266, RPMI8226, MM1S, and MOLP8 cells. Approximately 1 μCi of [^68^Ga]Ga-AJ206 was incubated with 1×10^6^ cells for 60 min at 4 °C. After incubation, cells were washed three times with ice-cold PBS containing 0.1% tween-20 and counted on an automated gamma counter (1282 Compugamma CS, Pharmacia/LKB Nuclear, Inc., Gaithersburg, MD). Blocking was performed with 2 μM of AJ206 to demonstrate CD38-specific cellular binding of [^68^Ga]Ga-AJ206. All cellular uptake studies were performed in quadruplicate for each cell line and repeated three times.

### Flow cytometry analysis of CD38 expression

To evaluate CD38 surface expression, cells (1x10^6^) were stained with PE-labelled anti-human CD38 monoclonal antibody (clone HB-7, BioLegend Cat # 356604) in FACS buffer (0.5% FBS with 2 mM EDTA) for 30 min on ice. After washing and resuspending in FACS buffer, data was acquired on a BD Accuri C6 plus flow cytometer and analyzed using FlowJo software. Receptor density measurements were performed using Quantibrite Beads (BD Biosciences, cat #340495), which contain four levels of phycoerythrin (PE) per bead. Gates were drawn on Low, Medium Low, Medium High, and High PE binding beads, and the geometric mean (FL-2 A: PE-A) from these populations was correlated with the lot-specific PE molecule/bead on a logarithmic scale. This correlation was used to translate cell population geometric mean to receptor/cell for the respective cell type.

### Western blotting

To assess CD38 protein expression in MM cells, lysed cells were denatured using Laemmli SDS sample buffer with beta-mercaptaethanol, loaded onto SDS-PAGE gels, and transferred to PVDF membranes. Blots were probed with rabbit monoclonal anti-human CD38 antibody (Clone RM388, Thermo Scientific, cat# MA5-36061) after blocking with 5% BSA. GAPDH was used as a loading control (Rabbit polyclonal, Clone D16H11, CST, cat# 5174). Blots were incubated overnight at 4°C, washed, and incubated with anti-rabbit IgG, HRP-linked secondary antibody (CST, cat #7074) to detect binding. Immunoreactivity was detected using super signal west Pico plus Chemiluminescent substrate (Thermo Scientific) and analyzed on iBright CL1500 Imaging System using Chemidoc scanner.

### Tumor models

Animal studies were performed under Johns Hopkins University ACUC-approved protocols. Male and Female NSG mice (5-6 weeks old) were used to establish xenografts by administering 3 million cells (50% Corning Matrigel) subcutaneously to form various tumor models within 3-4 weeks. Imaging or biodistribution studies (n=3-5) were conducted on mice with tumor volumes of 100-200 mm^3^. Disseminated tumor models were developed by injecting 4-5 million cells in saline intravenously for MOLP8 and MM1S-Luc models (n=4-5). Luciferase expressing MM1S cell line (MM1S-Luc) was used to monitor cancer cell dissemination in the body. Tumor establishment was confirmed by injecting D-Luciferin intraperitoneally (150 mg/kg) and imaging mice after 10 min in IVIS bioluminescence imaging system. Bone marrow tumors were ready for imaging after 25-30 days of implantation.

To establish PDX models, immune-deficient NSG male mice (NOD/Shi-scid IL-2rgnull), aged 6-8 weeks, were employed after being irradiated at an absorbed dose of 200 cGy. PCL cells, sourced from two patients and anonymized, were obtained from Dr. Gocke’s laboratory. Under sterile conditions, the cells were treated with DNAse at a concentration of 20μg/mL. After cell counting, 30,000 cells were subjected to use for flow cytometry analysis to assess CD38 expression. The remaining cells were subcutaneously inoculated at a concentration of 5×10^6^ cells in 100 μL of a solution comprising 60% Matrigel® Basement Matrix (Corning™ 356230) diluted with HBSS. The animals were included in the experiments once the PDX volumes reached tumor volumes of 100-200 mm^3^.

Bone marrow and peripheral blood samples were collected from MM patients or healthy donors who gave informed consent, in compliance with the Declaration of Helsinki. This was approved by the Institutional Review Board at the Johns Hopkins Medical Institutes. The mononuclear cells were then isolated via density centrifugation using Ficoll-Paque, a product of Pharmacia, located in Piscataway, NJ.

### Evaluation of pharmacokinetics of [^68^Ga]Ga-AJ206

Dynamic PET images were acquired on a Simultaneous 7T Bruker PET-MR scanner to evaluate the pharmacokinetics of [^68^Ga]Ga-AJ206. Mice bearing MOLP8 tumor xenografts were anesthetized under 2.5% isoflurane and a catheter was fixed in the tail vein before being secured on the PET-MR bed. An activity of ∼250 μCi (9.3 MBq) of [^68^Ga]Ga-AJ206 was administered intravenously and whole-body PET dynamic scans were performed starting from -1 to 5 min in 30-second intervals, followed by scans at 5-15 min in 1-minute intervals, 15-30 min in 3-minute intervals, 30-60 min in 5-minute intervals, and 60-90 min in 10-minute intervals. The acquired PET data were reconstructed and corrected for radioactive decay and dead time using ParaVision 360 V2 by Bruker. The percentage of injected dose per cc (%ID/cc) values were obtained by drawing ROI on the tumor, muscle, heart, liver and kidney using PMOD software, and image fusion and visualization were also performed using PMOD software.

### PET-CT imaging of mouse xenografts

Mice bearing flank tumors and disseminated disease were injected intravenously with [^68^Ga]Ga-AJ206 and PET images were acquired 60 minutes after injection of the radiotracer. Images were acquired in 2 bed positions for a total of 10 minutes using an ARGUS small-animal PET/CT scanner. Images were reconstructed using 2D-OSEM and corrected for radioactive decay and dead time. Image fusion, visualization, and 3D rendering were accomplished using Amira 2020.3.1. PET images were quantified by drawing ROI on tissues using Amide 1.0.6 and reported as %ID/cc.

### *Ex vivo* biodistribution

Mice with 100-200 mm^3^ tumor volume were used for *ex vivo* biodistribution studies. For radiotracer pharmacokinetics studies, mice with MOLP8 tumor xenograft received ∼80 μCi (2.96 MBq) [^68^Ga]Ga-AJ206 and were sacrificed at pre-determined time points (5, 30, 60 and 120 min). Similarly, for dosimetry study, mice with MM1S tumor xenografts received ∼80 μCi (2.96 MBq) of [^68^Ga]Ga-AJ206 and were sacrificed at pre-determined time points (5, 30, 60, 90 and 120 min). All other mice were injected with ∼50μCi (1.85 MBq) [^68^Ga]Ga-AJ206 and were sacrificed at 60 min. The selected tissues were collected, weighed, counted, and their %ID/g values calculated for biodistribution analysis. The tissues included blood, muscle, tumor, thymus, heart, lung, liver, pancreas, stomach, small intestine, large intestine, spleen, adrenals, kidney, bladder, femur, and brain.

### Harvesting of tissues for immunohistochemistry and flow cytometry

After the imaging study, mice were humanely euthanized and their tissues Including inflated lungs, and tumors were fixed in 4% paraformaldehyde and sent to Oncology tissue services (OTS) at JHU. Tissue sections of 4 μm thickness were prepared from paraffin-embedded tissues. Bones that were harvested were fixed and decalcified for 15 days in 10% EDTA on a slow-speed rocking shaker.

For flow cytometric analysis, single-cell suspensions were prepared from bone marrow and lungs. Femurs were properly blenched and cut from the epiphysis, and cells were collected directly in tubes. Lungs were chopped and digested in a buffer containing collagenase and DNase, incubated at 37°C for 30 minutes, and strained through a 40um cell strainer. Cell pellets were lysed in RBC lysis buffer, and live/dead staining was conducted using Near Infra-red 780 dye and CD38 staining, as previously described.

### ATRA treatment studies

To evaluate the pharmacodynamic effects of ATRA therapy on MM cells, U266, RPMI8226, MM1S, and MOLP8 cells were seeded at a concentration of 0.5 million in 500 μL per well in triplicates in a 6 well plate. A 200 nM working solution of All-Trans Retinoic Acid (ATRA, Sigma-Aldrich #554720) was prepared by diluting it in culture media. Next, 500 μL of the working solution was added to each well, and the cells were incubated for 48 hours at 37°C in a 5% CO2 atmosphere. After incubation, the cells were transferred to FACS tubes, the medium was removed by centrifugation, and the cells were resuspended in FACS buffer for flow cytometry and *In vitro* binding studies, following the same protocol as previously described.

To assess the pharmacodynamics of ATRA-induced CD38 expression *In vivo*, mice bearing PDX tumors (n=6) were subjected to image using [^68^Ga]Ga-AJ206 to quantify CD38 expression prior to treatment. Following the pre-treatment imaging, the mice were orally administered 500 μg of ATRA per day, prepared in a solution of 1% methyl cellulose, for a duration of 3 days. PET images were acquired on the fourth day after the start of treatment. Subsequently, the mice were humanely euthanized, and their tumors were collected in RPMI media for flow cytometry analysis and in formalin solution for IHC. All images were analyzed using AMIDE software with standard ROI analysis techniques.

### Immunohistochemistry

To perform immunohistochemistry, the tissue slides were deparaffinized by baking at 60°C and washing with xylene and alcohol. Antigen retrieval was carried out using citrate buffer (pH 6.0, 95-100°C, 20 min) and the endogenous peroxidase and alkaline phosphatase activity was blocked using BioXALL. The primary anti-human CD38 antibody was applied at a dilution of 1:250 and incubated overnight at 4°C (Thermo Scientific clone RM388 Cat# MA5-36061). After washing with PBS, the secondary antibody, Signalstain Boost IHC Detection Reagent (HRP), was applied and incubated for 30 min at room temperature. The slides were washed and developed using ImmPACT DAB substrate (Vector Lab #SK4105). After washing, the slides were counterstained with Mayer’s Hematoxylin for 1 min, dehydrated using alcohol and xylene, and then cover slipped.

### Statistical analysis

All statistical analyses were performed using Prism 9.0 Software (GraphPad Software, La Jolla, CA). Unpaired Student’s *t test* and one-or two-way ANOVA were utilized for column, multiple column, and grouped analyses, respectively. Paired student *t test* was used to analyze ATRA treatment studies. Statistical significance was set at ns, *P* ≥ 0.05; *, *P* ≤ 0.05; **, *P* ≤ 0.01; ***, *P* ≤ 0.001; ****, *P* ≤ 0.0001. Correlation was performed using simple linear regression without keeping the term constant at zero.

## Data availability

All study data are included in the article and/or supplementary information. Additional data are available upon request from the corresponding author.

## Authors’ contributions

**A.K. Sharma:** Conceptualization, methodology, data curation, formal analysis, writing–original draft, writing– review and editing. **K. Gupta:** Conceptualization, methodology, data curation, formal analysis, writing–original draft, writing–review and editing. **A. Mishra:** Methodology, data curation, writing–review and editing **G. Lofland:** Data curation, writing–review and editing **I. Marsh:** Data curation, writing–review and editing. **D. Kumar:** Data curation, writing–review and editing. **G. Ghiaur:** Methodology, formal analysis. **P. Imus**: Formal analysis, writing–review and editing. **S.P. Rowe:** Formal analysis, Writing–review and editing. **R.F. Hobbs:** Formal analysis, writing–review and editing. **C.B. Gocke**: Methodology, writing–review and editing. **S. Nimmagadda:** Funding acquisition, conceptualization, methodology, data curation, formal analysis, writing– original draft, writing–review and editing.

## Acknowledgements

This study was funded by NIH 1R01CA236616 (S. Nimmagadda) and the ^68^Ge/^68^Ga generator was supported by NIH R01CA269235 (S. Nimmagadda). Core resources (histology and imaging) were supported by NIH P30CA006973. We thank Drs. Aykut Üren and Purushottam Tiwari at Georgetown University for performing SPR experiments.

