## Supplementary information for "A Gallium-68-Labeled Peptide Radiotracer For CD38-Targeted Imaging In Multiple Myeloma With PET"

for

**Running title:** New Radiotracer for CD38-Targeted Imaging in Multiple Myeloma

**Key words:** CD38, PET, Daratumumab, Isatuximab, Minimal residual disease

**Funding:** This study was funded by NIH 1R01CA236616 (SN) and the Ga-68 generator was supported by NIH R01CA269235. Core resources (histology and imaging) were supported by NIH P30CA006973.

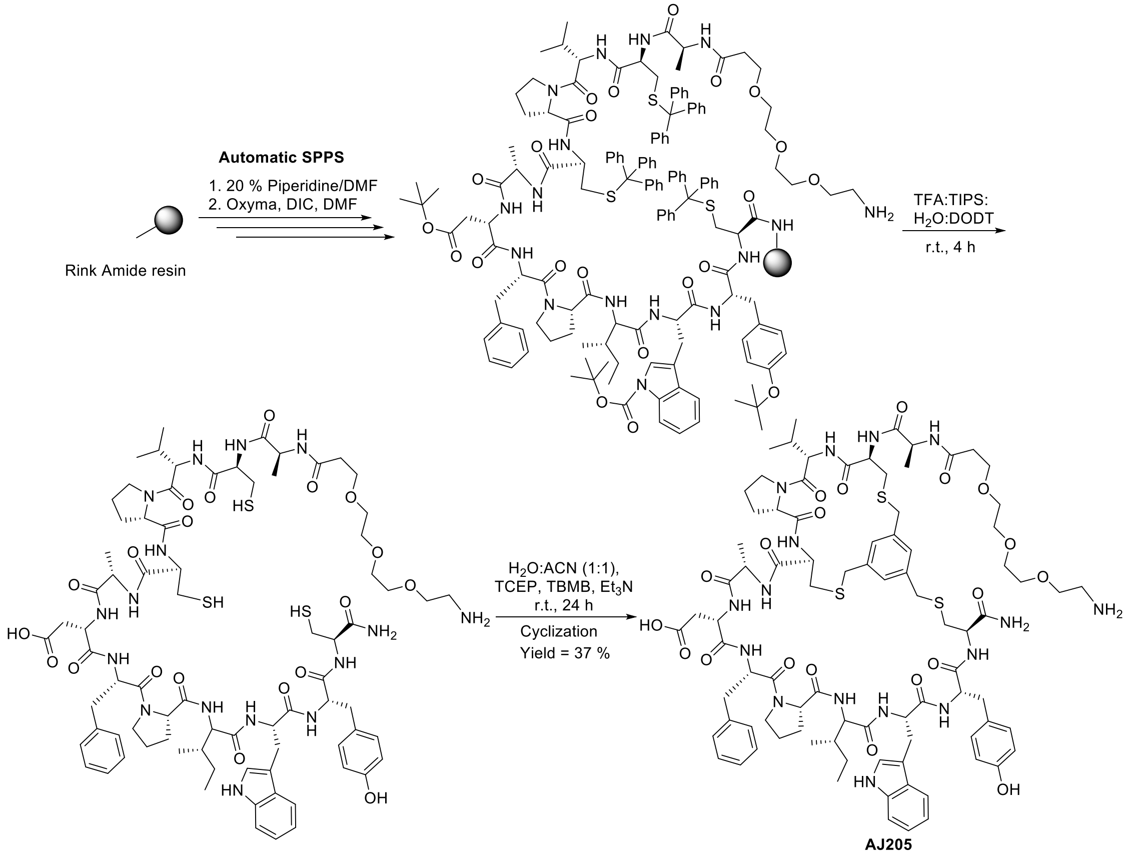


**AJ205**
**$NH_2$-PEG$_3$-(ACVPCADFPIWYC-$NH_2$)TBMB**

**Supplementary scheme 1**. Synthesis of AJ205**.** Automatic solid phase peptide synthesis by adding Fmoc-protected amino acids to Rink amide resin using microwave assisted coupling reaction. The created peptidyl resin was treated with cleavage cocktail to obtain deprotected linear peptide. Liner peptide was cyclized with TBMB in the presence of $Et_3N$ in water:acetonitrile mixture to obtain AJ205.

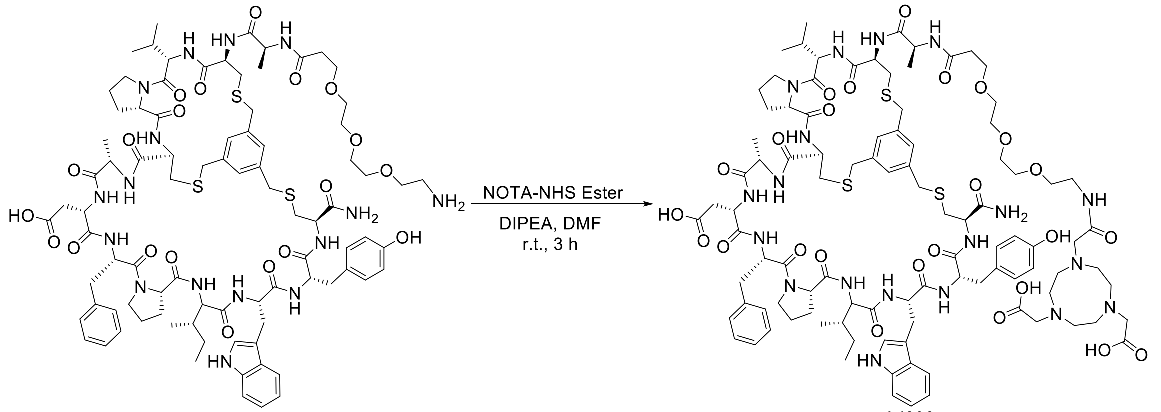


AJ205

AJ206
NOTA-PEG$_3$-(ACVPCADFPIWYC-$NH_2$)TBMB

**Supplementary scheme 2.** Conjugation of AJ205 with NOTA-NHS ester in the presence of DIPEA to obtain AJ206

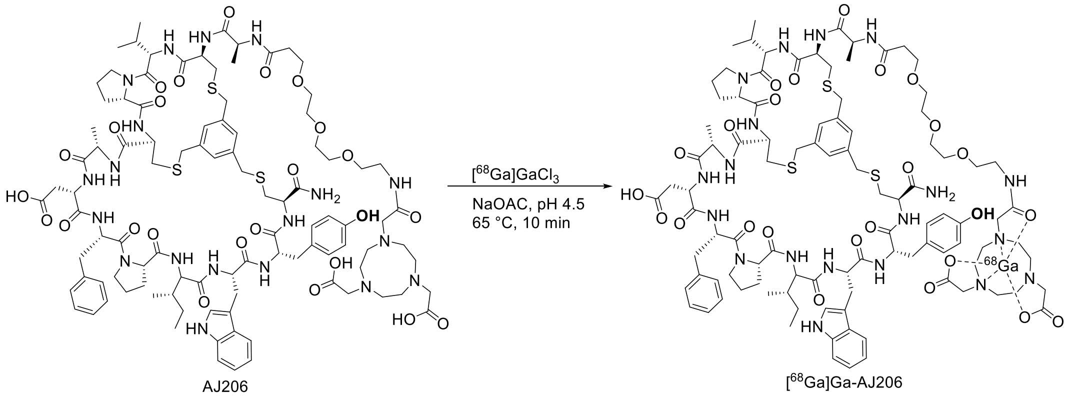


**Supplementary scheme 3.** Radiolabeling reaction of AJ206 with [$^{68}$Ga]GaCl$_3$

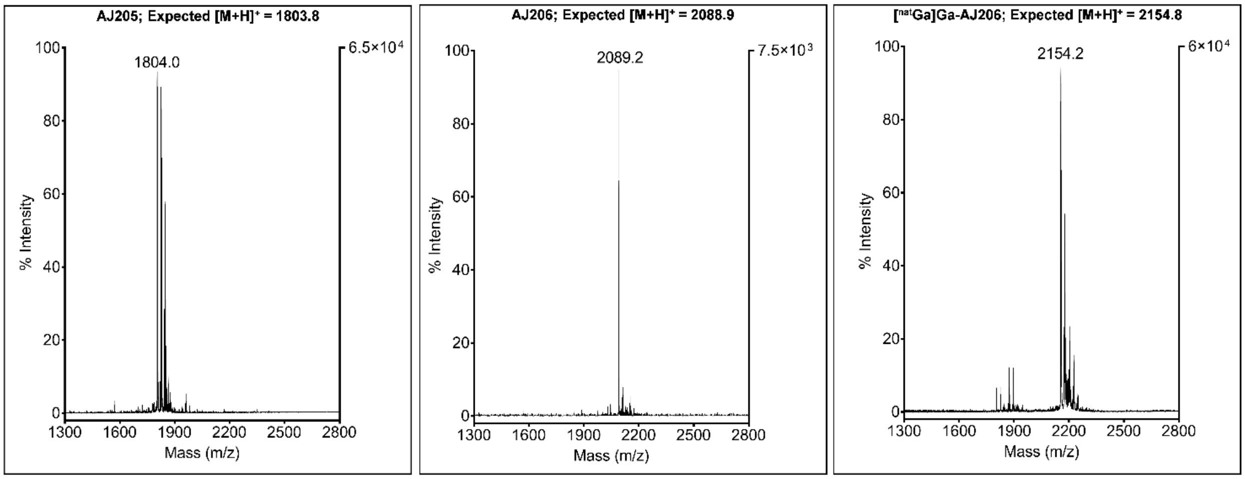


**Supplementary figure 1.** Characterization of AJ205, AJ206 and [$^{nat}$Ga]Ga-AJ206 using MALDI-TOF Mass spectrometry

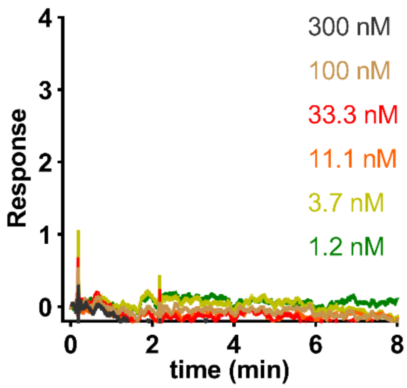


**Supplementary figure 2.** Surface plasmon resonance (SPR) analysis of AJ206 with purified recombinant mouse CD38 protein showing no binding to mouse CD38.

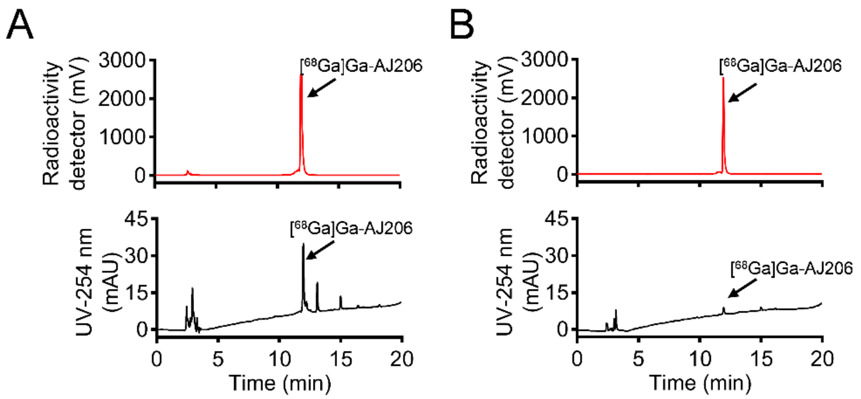


Decay corrected Radiochemical yield (RCY) = 92 ± 10.5 % (n = 35)

Radiochemical purity (RCP) = 98 ± 0.5 % (n = 35)

Specific activity = 10-15 GBq/µmol (270-400 mCi/ µmol)

**Supplementary figure 3.** Radiolabeling and characterization of [$^{68}$Ga]Ga-AJ206. **A)** HPLC chromatograms of reaction mixture. **B)** HPLC chromatogram of purified radiolabeled product after formulation.

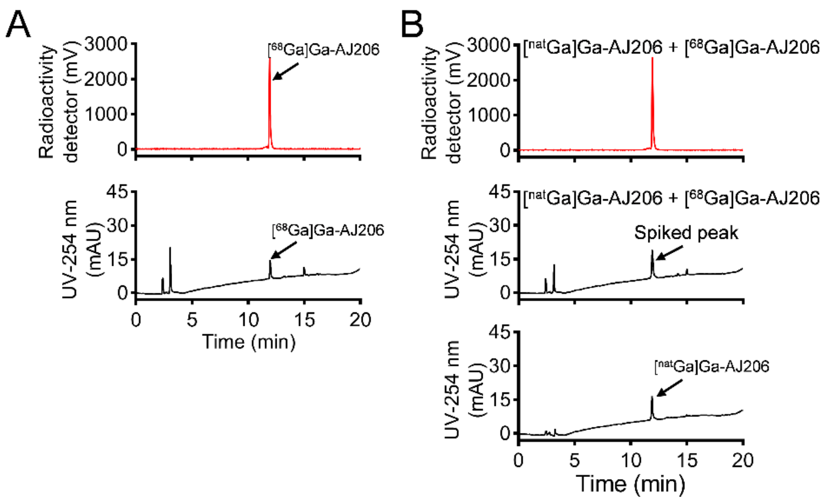


**Supplementary figure 4.** Characterization of [$^{68}$Ga]Ga-AJ206. **A)** Stability of radiolabeled product in formulation buffer at 120 min. **B)** HPLC chromatogram of chemical identity of [$^{68}$Ga]AJ206 with [$^{nat}$Ga]Ga-AJ206

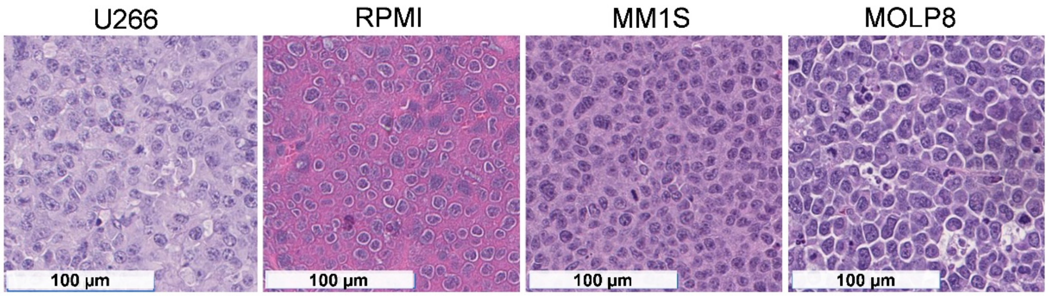


**Supplementary figure 5.** H&E stained slides of various MM tumor xenograft models

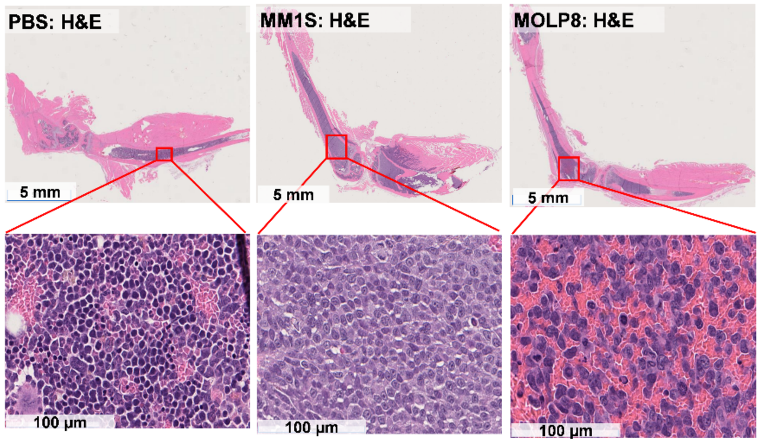


**Supplementary figure 6.** H&E stained slides of bone section of NSG mice intravenously injected with PBS, MM1S and MOLP8 cells

**Supplementary table 1.** Biodistribution of [$^{68}$Ga]Ga-AJ206 in mice with MM1S xenografts; data is presented as Mean±SEM (n=4) of %ID/g.

| | **MM1S tumor xenograft at different time-points (Mean±SEM)** | | | | |
|---|---|---|---|---|---|
| **Tissues** | **5 min** | **30 min** | **60 min** | **90 min** | **120 min** |
| **Blood** | 20.66±1.68 | 2.95±0.37 | 0.75±0.29 | 0.75±0.24 | 0.33±0.07 |
| **Muscle** | 1.96±0.28 | 0.55±0.09 | 0.13±0.02 | 0.18±0.04 | 0.09±0.02 |
| **Tumor** | 3.09±0.16 | 4.15±0.53 | 2.48±0.17 | 1.60±0.46 | 1.30±0.40 |
| **Thymus** | 16.54±5.34 | 2.67±0.83 | 0.69±0.19 | 0.84±0.18 | 0.33±0.10 |
| **Heart** | 6.38±0.42 | 1.00±0.15 | 0.29±0.03 | 0.35±0.08 | 0.16±0.01 |
| **Lung** | 10.18±4.52 | 3.43±0.50 | 0.67±0.07 | 0.95±0.18 | 0.40±0.04 |
| **Liver** | 30.31±3.51 | 3.64±0.31 | 1.38±0.04 | 1.16±0.08 | 0.73±0.10 |
| **Spleen** | 6.41±1.30 | 1.21±0.15 | 0.41±0.05 | 0.43±0.07 | 0.28±0.05 |
| **Pancreas** | 3.55±0.34 | 0.79±0.14 | 0.19±0.03 | 0.26±0.06 | 0.17±0.05 |
| **Adrenals** | 5.07±0.11 | 2.08±0.38 | 0.59±0.06 | 0.44±0.10 | 0.57±0.14 |
| **Kidney** | 31.50±4.15 | 21.65±2.02 | 14.09±0.57 | 13.24±1.59 | 9.39±1.50 |
| **Ovary** | 4.11±1.40 | 1.48±0.32 | 0.20±0.06 | 0.32±0.14 | 0.45±0.15 |
| **Bladder** | 9.35±1.87 | 4.51±0.93 | 0.75±0.08 | 1.23±0.10 | 0.90±0.37 |
| **Stomach (with contents)** | 1.65±0.19 | 0.54±0.19 | 0.32±0.21 | 0.82±0.36 | 0.47±0.30 |
| **Small intestine (with contents)** | 5.00±0.70 | 0.95±0.10 | 0.20±0.10 | 0.41±0.08 | 0.52±0.21 |
| **Large intestine (with contents)** | 3.11±0.38 | 0.75±0.04 | 0.29±0.03 | 0.27±0.05 | 0.27±0.19 |
| **Femur** | 3.37±0.31 | 0.86±0.06 | 0.33±0.12 | 0.28±0.05 | 0.09±0.02 |
| **Brain** | 0.44±0.07 | 0.21±0.12 | 0.06±0.03 | 0.03±0.01 | 0.02±0.01 |
